# Ktd1 is a phospho-regulated member of the Dup240 family that mediates defence against killer toxin K28

**DOI:** 10.64898/2026.08.07.743514

**Authors:** Hatwan Nadir, Alex Pembery, Kamilla Laidlaw, Amy Milburn, Meru Sadhu, Mark C. Leake, Chris MacDonald

## Abstract

The *DUP240* gene family in *Saccharomyces cerevisiae* encodes ten proteins containing two transmembrane domains (TMDs). Despite decades of interest driven by their high sequence similarity, little functional information exists regarding whether Dup240 family members share redundant or distinct roles. In this study, we combined computational modelling, subcellular localisation, and functional assays across the family to identify shared and unique features. Computational modelling revealed that the Dup240 member Ktd1 possesses a unique structural element adjacent to its TMD region. Out of six successfully localised family members, Ktd1 was the only protein predominantly targeted to the vacuolar membrane and the sole Dup240 required for defence against the K28 killer toxin. Computational predictions further indicated that Ktd1 undergoes extensive post-translational regulation, containing multiple validated phosphorylation sites. Screening potential regulatory kinases and phosphatases identified several enzymes required for K28 defence, which were independently validated using liquid-based toxin sensitivity assays. A multicopy suppressor screen demonstrated that *KTD1* overexpression rescued K28 sensitivity across most enzyme mutant backgrounds, confirming Ktd1 acts downstream or in parallel to many factors. However, the phosphatase Sit4 and the kinase Hog1 scored as most likely co-factors in Ktd1 mediated defence. Live-cell fluorescence imaging of these two enzymes revealed no dramatic spatial re-localisation during K28 exposure, suggesting that phospho-dependent regulation of Ktd1-mediated defence may occur through transient signalling events. Together, these findings identify Ktd1 as the central effector of the Dup240 family in toxin defence and provide a mechanistic framework for understanding Dup240 regulation.

## INTRODUCTION

Protein toxins are weapons secreted by cells in a hostile battle against other organisms (Aktories and Schmidt, 2015). The A/B family of toxins includes notable includes notable examples known to impact human health, such as cholera toxin, diphtheria toxin, Shiga toxin, anthrax toxin, and ricin (Odumosu et al., 2010). A/B toxins are defined by a bipartite structure of active (A) and binding (B) sections and manipulate host cell trafficking pathways after gaining access to the cell by binding to sugars and entering through endocytosis (Beddoe et al., 2010; Márquez-López and Fanarraga, 2023). The dissociation of the A and B parts allows for targeted attack on organelles, such as causing cell cycle arrest (Jinadasa et al., 2011; Teter, 2013). One example of an A/B toxin is that of the virally encoded killer toxin 28 (K28), which is secreted by infected yeast to kill other susceptible yeast (Becker and Schmitt, 2017; Schmitt and Breinig, 2006). Ktd1 is a membrane protein encoded by the *DUP240* gene family and was discovered as a rapidly evolving defence factor against K28 (Andreev et al., 2023).

The genome sequencing project in *S. cerevisiae* first identified members of the *DUP240* gene family, including *DFP3* on Chromosome III and six homologues on chromosome I, five of which are arrayed in tandem (Bussey et al., 1995; Goffeau et al., 1996). The first comprehensive bioinformatic and functional mapping of the *DUP240* family revealed that it comprises ten paralogues encoding ∼240-amino-acid, membrane-associated proteins (Poirey et al., 2002). In the S288C reference strain, seven members are clustered in tandem arrays on Chromosomes I and VII, while the remaining three exist as singletons on Chromosomes I, III, and VIII. These loci exhibit high genomic instability and dynamic copy-number variation across diverse yeast strains. Owing to the nature of these tandem arrays, the total count of *DUP240* genes fluctuates dramatically between strains, demonstrating diversifying selection via rapid evolution (Andreev et al., 2023; Leh-Louis et al., 2004) Subsequent evolutionary analyses expanded this framework by defining the overarching *DUP* gene family, which encompasses 23 total members divided into the *DUP240* subfamily and the larger *DUP380* subfamily arising from a domain duplication event, which encode tetraspan membrane proteins of ∼380-amino-acids (Despons et al., 2006). While *DUP380* genes are concentrated in subtelomeric loci of different chromosomes in the S288C genome, subsequent genetics revealed they form dynamic tandem arrays in some other yeast strains, driving further chromosomal plasticity (Despons et al., 2011).

The *DUP380* genes encode the Cos proteins, a functionally redundant family of endolysosomal proteins that are metabolically regulated by extracellular NAD^+^ levels sensed by the sirtuin, Sir2 (MacDonald et al., 2015a). Cos proteins serve as adaptors to facilitate lysosomal trafficking of various cargoes, including examples like GPI-anchored proteins that rely entirely on Cos proteins to access the ESCRT-mediated multivesicular body pathway for lysosomal degradation (MacDonald et al., 2015b). The function of the Dup240 proteins has been more elusive, but some work has revealed aspects of their biology. Targeted deletion of all ten paralogues simultaneously does not impair cell growth, showing the Dup240 family is non-essential for viability (Poirey et al., 2002). Subcellular localisations from this study confirmed the prediction that Dup240 proteins contain two transmembrane domains and anchor to membranes within the yeast system. *PRM8* and *PRM9* genes were shown to encode membrane proteins that are upregulated in response to mating factor, but this role remains undefined (Heiman and Walter, 2000). A potential role in retrograde protein transport from the Golgi to endoplasmic reticulum (ER) vesicles has been proposed as *MST27* was identified from a multicopy suppressor screen for COPI mutant subunit *sec21-3*. Both Mst27 and its paralogue Mst28 have C-termini motifs that can bind COPI and COPII subunits (Sandmann et al., 2003).

More recently, quantitative trait locus (QTL) mapping of yeast strains with varied sensitivity to the killer toxin K28 identified a narrow genomic interval harbouring only two genes: *UIP3* and *KTD1*, both encoding Dup240s (Andreev et al., 2023). Although no evidence for a role in K28 defence was revealed for Uip3, Ktd1 was shown to be required for tolerating K28 toxin. Further work showed that trafficking of Ktd1 through the endolysosomal system is required for K28 defence (Laidlaw et al., 2025); whether additional co-factors or posttranslational regulators contribute to this process is currently unknown. Additionally, a potential role for other Dup240 family members contributing to Ktd1-mediated toxin defence has not been systematically tested. In this study we performed a computational comparative analysis of the Dup240 protein sequences, predicting structures and likelihood of their regulation by post-translational modifications. Our *in vivo* functional experiments went on to show that Ktd1 has a specific and unique localisation to the vacuole membrane, and specific phosphorylation sites yet unidentified in other Dup240s are required for K28 defence. This prompted us to screen for phospho-regulated genes across the yeast genome for roles in K28 defence using Ktd1 suppression to prioritise likely regulators of the Ktd1 pathway.

## RESULTS

### Comparative analysis of the Dup240 family

Structural predictions were mapped for each Dup240 protein using AlphaFold3 (Abramson et al., 2024), with predicted local distance difference test (pLDDT) confidence presented as a heat map (**Sup. Figure S1A - S1B**). As expected within a family of related proteins, the predicted structures of Dup240 members share many features (**Figure 1A**). *DFP2* was excluded from this analysis as in S288C it encodes a 74 amino acid protein lacking trans-membrane domains; it appears a nonsense mutation truncates the canonical Dup240 prematurely, as its non-coding 3’UTR sequence shares homology with the family. We also excluded 14 more recently identified homologues in other *S. cerevisiae* strains (Andreev et al., 2023). Dfp1 was included despite the canonical start codon being skipped, as the remaining Dup240 elements from the trans-membrane domain (TMD) onwards is retained (Wirth et al., 2005). The N-termini and first transmembrane domains of Dup240 proteins have high levels of sequence homology, while the luminal facing residues and the second TMD display most sequence divergence (**Figure 1B - 1D, S1C**). Previous analysis of Ktd1 and its homologues revealed specific codons in these regions under positive evolutionary selection, suggesting rapidly evolving amino acid sites that might be important for toxin defence (Andreev et al., 2023). Intriguingly, Ktd1 has been proposed to form ring-like oligomers (Wang et al., 2023), and we find that the rapidly evolving residues are strongly enriched towards facing the interior of the proposed ring (**Sup. Figure 2**). The luminal region that is important for Ktd1 function contains a cysteine residue, as do five other Dup240s (**Figure 1G**), which is not predicted to form an intramolecular disulfide bond, but could be available for intermolecular interactions. We note that Ktd1 has 1 unique predicted coiled domain close to the TMD region (**Sup. Figure S1D**). The C-termini have a mixture of regions that are not well conserved across Dup240s, but also specific regions/motifs (**Figure 1E - 1F**). One example is a YYFY at residue 154 in consensus sequence, which may facilitate an additional membrane interaction as a hydrophobic anchor with flat, aromatic rings inserted directly into the inner leaflet (Muhammedkutty and Zhou, 2026). There are also an asparagine / glutamine NQ pairs (consensus residue 142) and a conserved lysine (consensus residue 220) that is represented in all 9 selected Dup240s. The latter is potentially important for posttranslational modifications, such as ubiquitination or acetylation (Hao et al., 2024).

**Figure 1:**
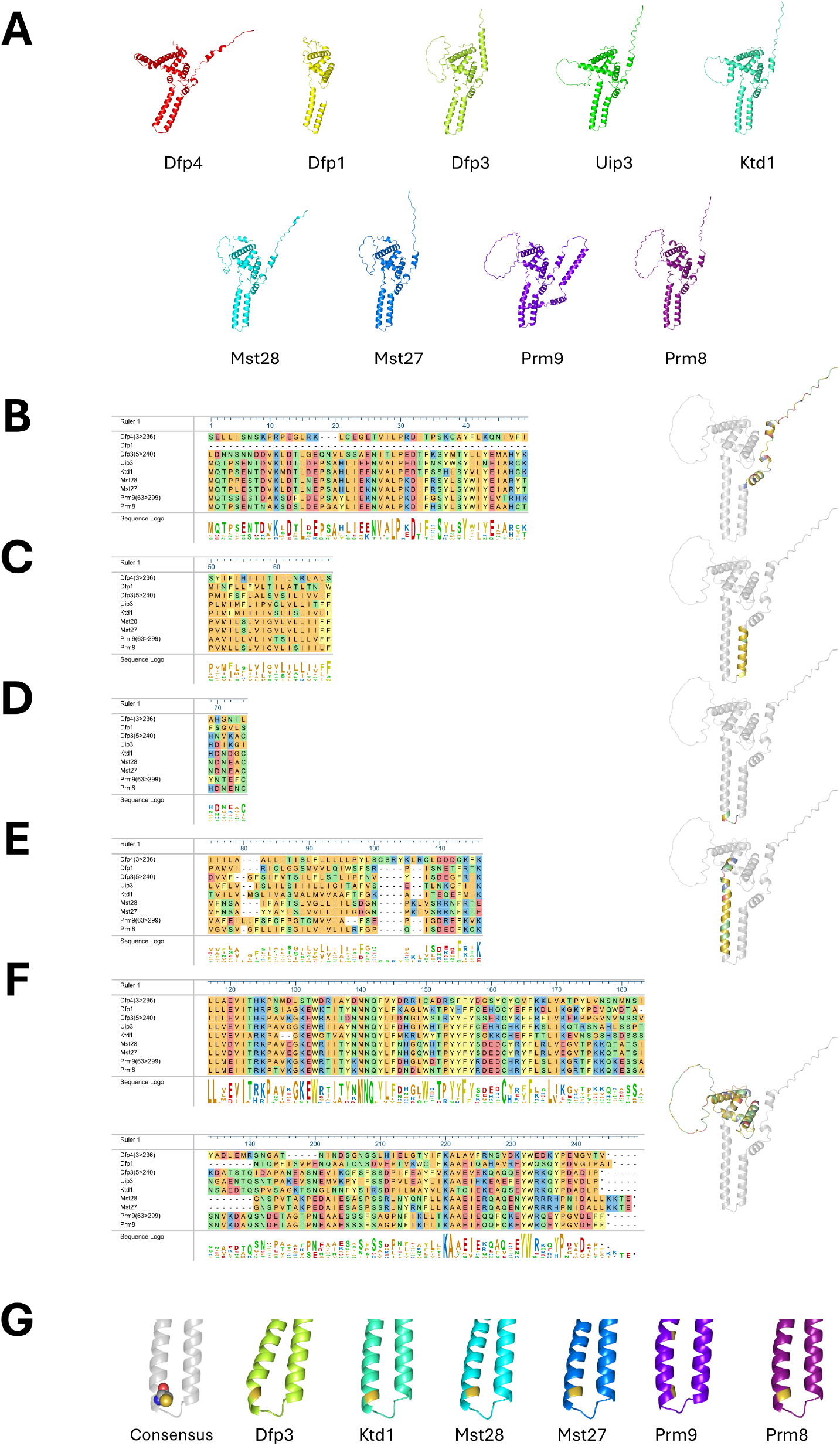
Comparative analysis of sequences and predicted structures of the Dup240 family. **A)** AlphaFold3 predicted structures for nine of the ten Dup240s found in *S. cerevisiae*. **B-F)** MUSCLE sequential alignments of the Dup240s, separated by region. The section represented in each row is highlighted on a predicted AlphaFold3 structure of the consensus sequence of the Dup240 family (grey, right). Colours are defined by MegAlign Pro’s ‘Colour by Chemistry’ scheme. The sections are: cytoplasmic N-termini **(B)**, first transmembrane domain **(C)**, luminal linker region **(D)**, second transmembrane domain, with additional alpha helix **(E)**, cytoplasmic C-termini **(F). G)** Zoomed-in section of transmembrane domains joined by the luminal linker, with cysteines labelled in yellow on the cartoon representation. A structure of the Dup240 consensus sequence (grey) shows the conserved cysteine in a spherical representation.

**Figure 2:**
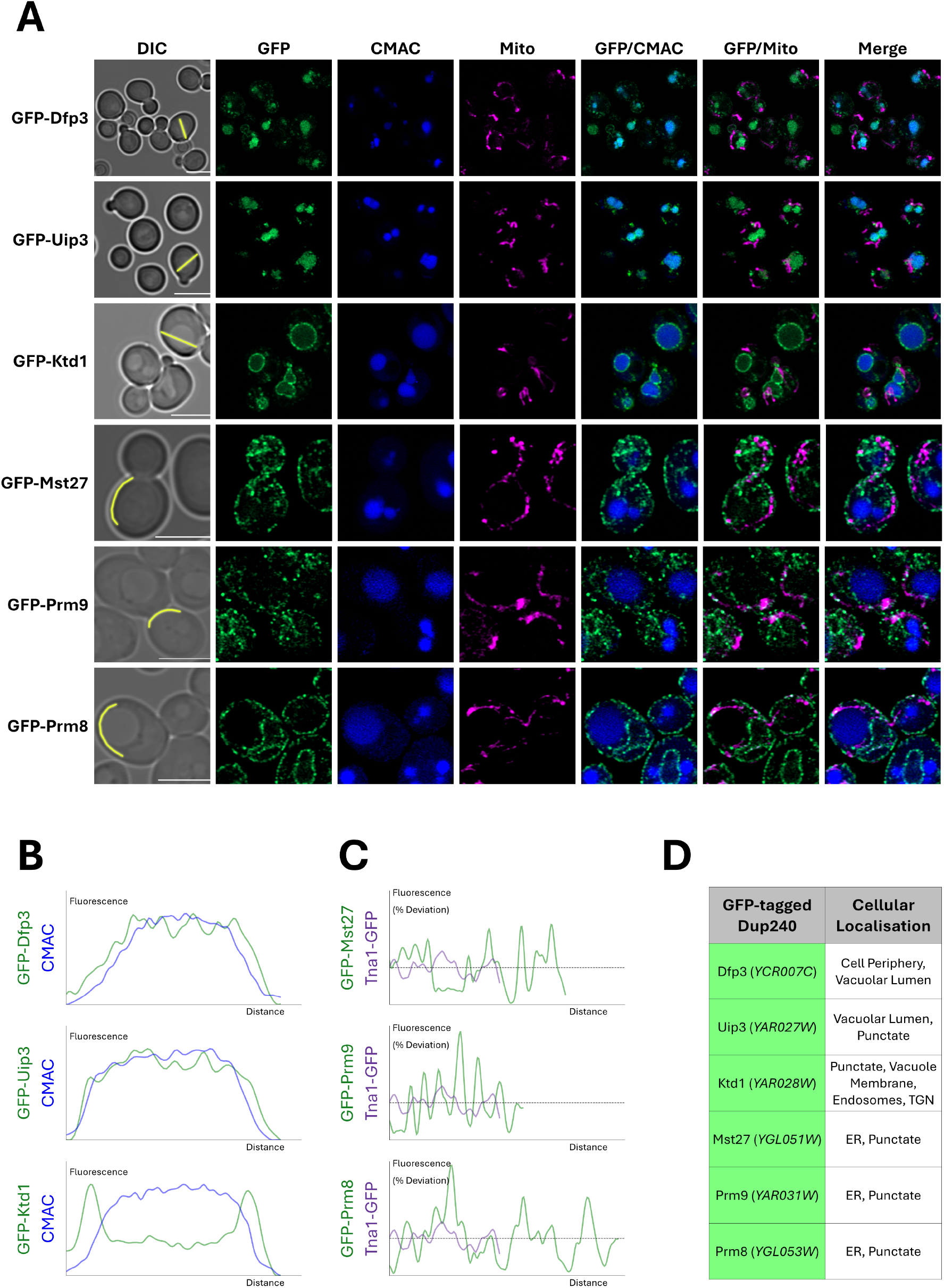
The Dup240 family exhibit distinct localisation patterns. **A)** Airyscan2 confocal micrographs of indicated GFP tagged Dup240 proteins, with CMAC (vacuole, blue) and MitoTracker Red (Mitochondria, magenta) labelling performed prior to imaging. **B -C)** Line analysis of normalised fluorescence (region shown in yellow, **(A)**) of GFP-Dup240s compared to the CMAC labelled vacuolar lumen **(B)** or the percentage deviations from the average: compared between line profiles of ER localised Dup240s (green) compared to a ER marker control, Tna1-GFP (purple) **(C). D)** Table outlining the shown Dup240 proteins, systematic gene names, and their observed localisations. Scale bar, 5μm.

Previous work has localised various Dup240 proteins using a C-terminal fluorescent tag approach to different cellular locations, including the plasma membrane (PM) and cytoplasmic puncta (Poirey et al., 2002). We previously compared various Dup240 tagging strategies to visualise Ktd1 and found an N-terminal GFP tag, expressed under control of the constitutive *NOP1* promoter, was optimal for localising Ktd1 to different endolysosomal compartments whilst also preserving protein function (Andreev et al., 2023). We applied this approach to other Dup240 members, followed by high resolution Airyscan2 imaging of cells labelled with CMAC for the vacuolar lumen and MitoTracker Red CMXros for the mitochondria. Of the six successfully localised Dup240s, we do not observe any at the plasma membrane, with family members observed at either the vacuole or the ER (**Figure 2A**). Both Dfp3 and Uip3 localise to the vacuolar lumen, with significant overlapping signal with CMAC (**Figure 2B**). The three other Dup240s successfully imaged, localised to the perinuclear and cortical ER. Previous work showed Mst27/28 and Prm8/9 have COP-binding motifs, CCXY and FFLL respectively (Sandmann et al., 2003), supporting this ER localisation phenotype. Strikingly, GFP-Mst27, GFP-Prm8 and GFP-Prm9 have a punctate phenotype within the ER membrane (**Figure 2C**) that is very distinct from classic ER localisation phenotypes, such as tagged versions of Sec63 and Tna1 (Paine et al., 2021), that exhibit a more uniform membrane localisation pattern (**Sup. Figure S3**).

### Toxin defence via Ktd1 is post-translationally regulated

Next, we tested the capacity for different *DUP240* deletion strains to protect against K28 toxin. Previously optimised halo assays were used to assess sensitivity, including precise media and culturing conditions (Laidlaw et al., 2025), coupled to an unbiased analysis pipeline. As previously reported, *ktd1Δ* cells are hypersensitive to K28, but other *DUP240* deletions had marginal effects (**Figure 3A**). These results were confirmed using a liquid based K28 assay, where null strains were grown in the presence on media enriched with K28 toxin, or a heat cured control, which showed *ktd1Δ*, but no other *DUP240* mutants, were hypersensitive (**Figure 3B**). We therefore conclude that Ktd1 is unique across the Dup240 family in its ability to mediate defence against K28 toxin. Features that might explain this specificity include its unique localisation to the limiting membrane of the vacuole, in addition to other endolysosomal compartments. Ktd1 also exhibits specific predicted structural features, like a membrane adjacent coiled-coil region.

**Figure 3:**
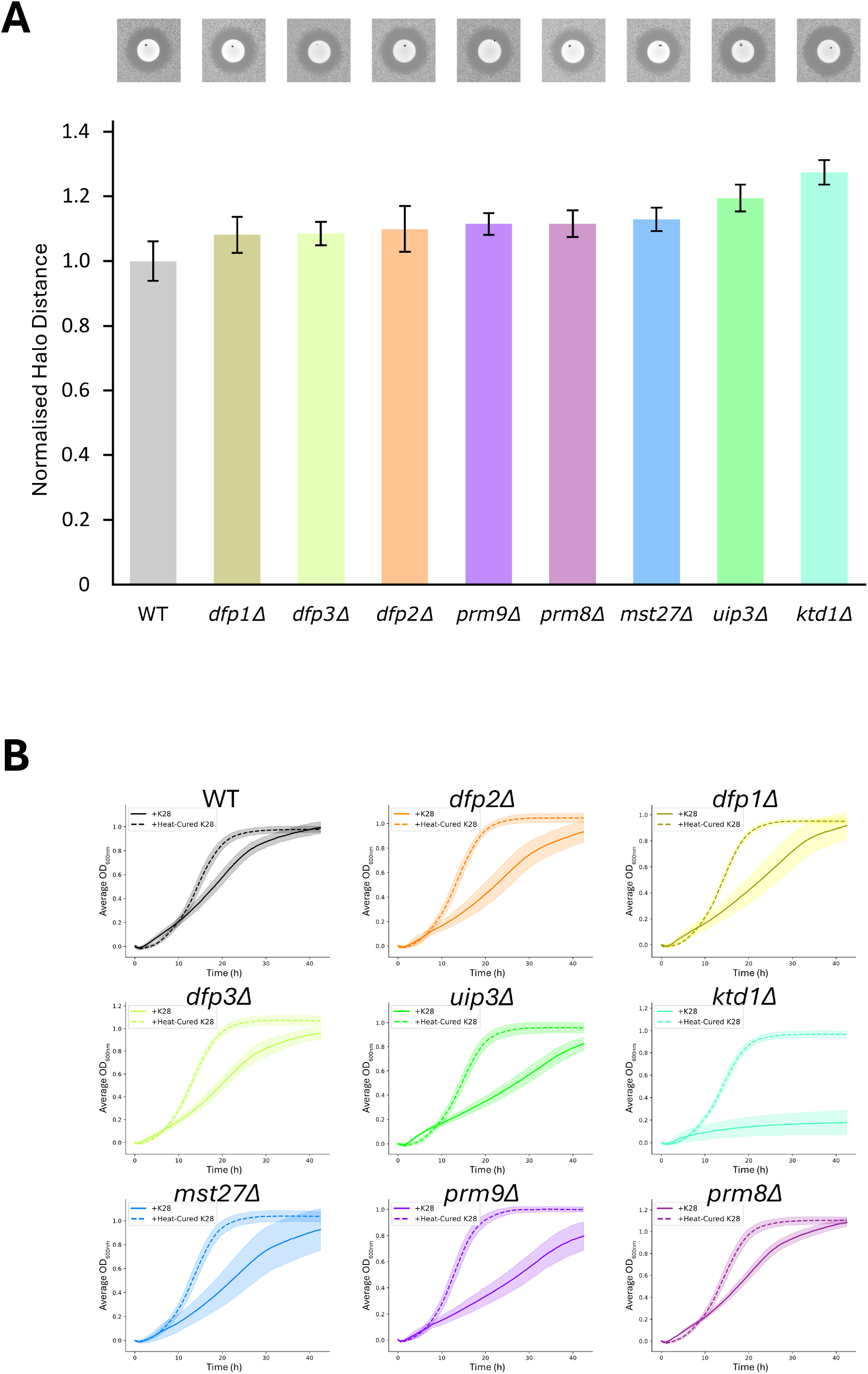
Ktd1 is the only member of the Dup240 family required for K28 resistance. **A)** Halo radii calculated by custom analysis pipeline of the *dup240Δ* cells plated on SC media pH 4.7 and exposed to spots of K28 secretor yeast from a separate culture. Values were normalised relative to wild type with the standard deviation indicated with error bars (n=4). A representative image of each assay is shown above. **B)** Liquid based growth assays of *dup240Δ* cells grown in the presence of K28 media (solid line) or an inactive heat cured control (dashed line) toxin, with the standard deviation plotted for each curve (shaded area).

Having confirmed and computationally highlighted features of Ktd1 and other Dup240s, we next employed complementary bioinformatic approaches to investigate how Ktd1 is regulated post-translationally. Analysis of potential phosphorylation sites, using NetPhosYeast (Ingrell et al., 2007), revealed a broad spectrum of candidate modifications, clustered predominantly within the N-terminal region and a predicted unstructured loop downstream of the second transmembrane domain (**Figure 4A**). The N-terminal predictions were conserved across several Dup240 proteins, including Ktd1, as was the cluster of serine residues towards the C-terminus, which also contained several Ktd1-specific sites (**Figure 4B - 4C**). These predictions are in agreement with mass spectrometry analyses showing Ktd1 residues T3, S5, T8 (denoted as cluster 1) and S170, S174, S176, S182, S185 (denoted as cluster 2) can be phosphorylated (Swaney et al., 2013; Zhou et al., 2021). Phospho-null mutations were targeted by mutating phosphorylated residues in each cluster to alanine (**Figure 4D**). Expression of the phospho-ablative mutants of cluster 1 (Ktd1^3A^) or cluster 2 (Ktd1^5A^) were unable to rescue the hypersensitive phenotype of *ktd1Δ* cells, unlike a control plasmid expressing wild-type Ktd1^WT^ (**Figure 4E - 4F**). This work shows that all Dup240 proteins are likely phosphorylated and in the case of Ktd1 at least, this phosphorylation is required for defence against K28. It may be that other Dup240s localised to distinct membranes utilise phosphorylation for analogous functions.

**Figure 4:**
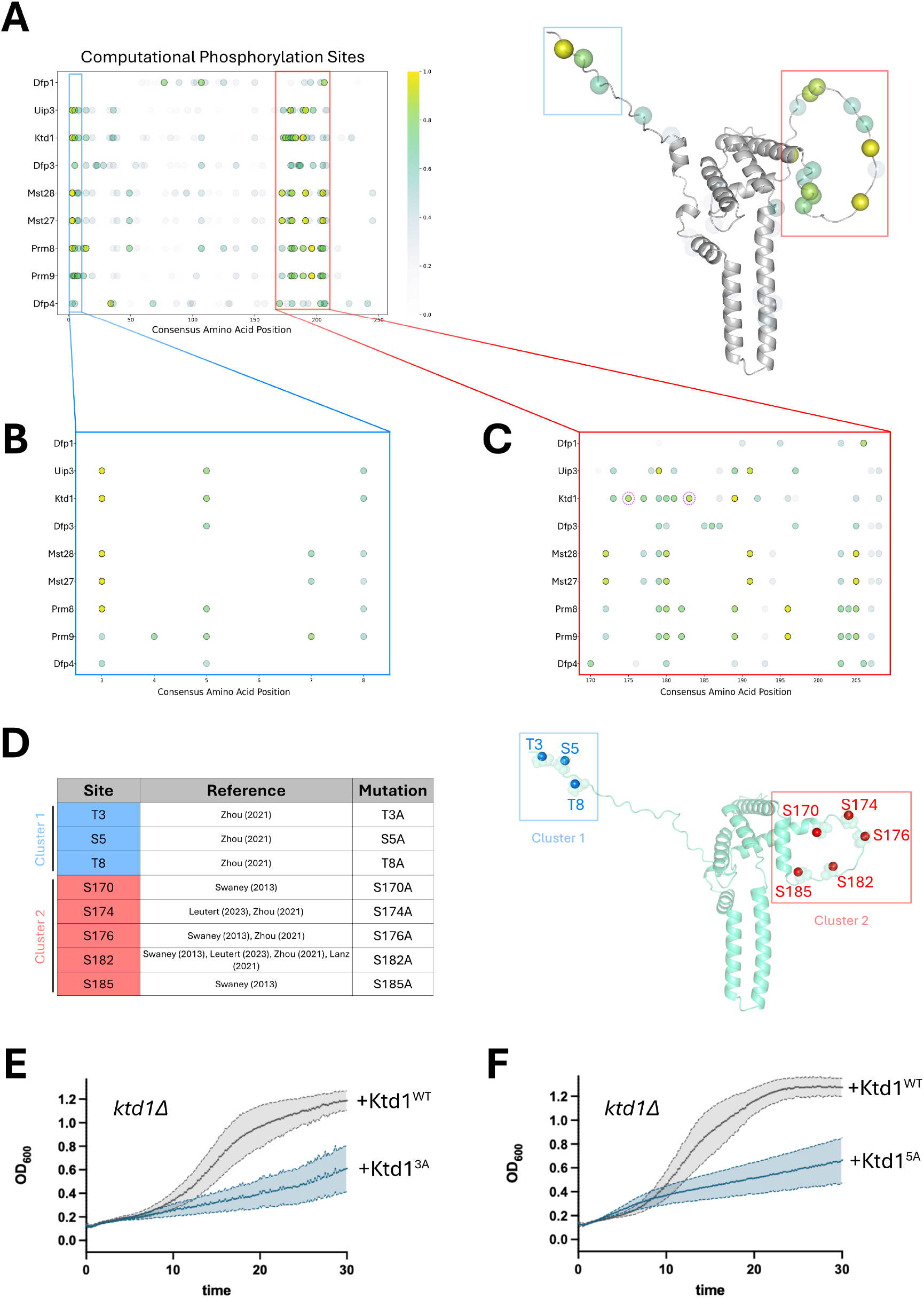
Ktd1 requires phosphorylation to mediate K28 defence. **A)** Computationally predicted phosphorylation sites for Dup240 proteins generated using NetPhosYeast. Confidence levels are shown by both the viridis colormap and colour transparency. An AlphaFold3 predicted structure for the Dup240 consensus provides a scaffold for display. **B - C)** Zoom of cluster 1 (residues 3 – 8), and cluster 2 (residues 170 – 208), of the Dup240 protein consensus. **D)** Observed phosphorylated serine and threonine residues in Ktd1 are listed (left) and mapped on to the predicted structure of Ktd1 (right). Cluster 1 (blue) includes 3 phosphosites and cluster 2 (red) includes 5 residues that are phosphorylated. **E - F**) Liquid growth assay of *ktd1Δ* cells transformed with a plasmid expressing wild-type Ktd1 (Ktd1^WT^) shown in grey or phospho-ablative mutations (blue) in cluster 1 (Ktd1^3A^, **E**) or cluster 2 (Ktd1^5A^, **F**).

### A genetic screen for phospho-regulators of Ktd1

Although a high throughput screen has been previously performed for K28 sensitivity in *S. cerevisiae* (Carroll et al., 2009), we have optimised assay sensitivity conditions to reveal more subtle mutations that were not identified in the original screen, such as the COG complex subunits (Laidlaw et al., 2025). Therefore, we arrayed a series of phosphatase and kinase mutants representing the enzymome that regulates phosphorylation in yeast. For this, deletion mutants of viable nulls were used for most genes (Giaever et al., 2002), but DAmP depletion mutants (Breslow et al., 2008) were used for any genes encoding essential enzymes (**Figure 5A**). Our optimised assays were used to screen for K28 sensitivity following exposure, using a high throughout analysis pipeline (Pembery et al., 2026). All normalised results for halo assays, including standard deviations, are included (**Sup. Table 1**). Of the 52 phosphatase mutants, the vast majority had very little phenotypic difference to K28 exposure when compared to wild-type cells (**Figure 5B**). Several mutants gave subtle resistance phenotypes, with halo distances (measured from the edge of secretor cell spot to the start of the lawn) as low as 0.7 ± 0.1 relative to wild-type cells, and the four most sensitive mutants encoded Sit1, Glc7, Ptc4 and Yvh1, with halo distances ranging from 1.24 ± 0.08 to 1.6 ± 0.1 (**Figure 5C**). The screen of 121 kinase related factors also gave a large percentage of mutants with very similar K28 sensitivity as wild-type cells. However, there was a larger number of kinase mutants with hypersensitive phenotypes, including several mutants with extremely high halo distances exceeding that of *ktd1Δ* mutants (**Figure 5D**). The most extreme examples were *bud23Δ* and *ctk3Δ*, which produced halo distances >3.0 ± 0.1. Six of these phosphorylation mutants have been previously identified from a genome-wide screen (Carroll et al., 2009), and we have implicated 26 novel candidates using the same sensitivity thresholding (**Sup. Figure S4A - S4B**).

**Figure 5:**
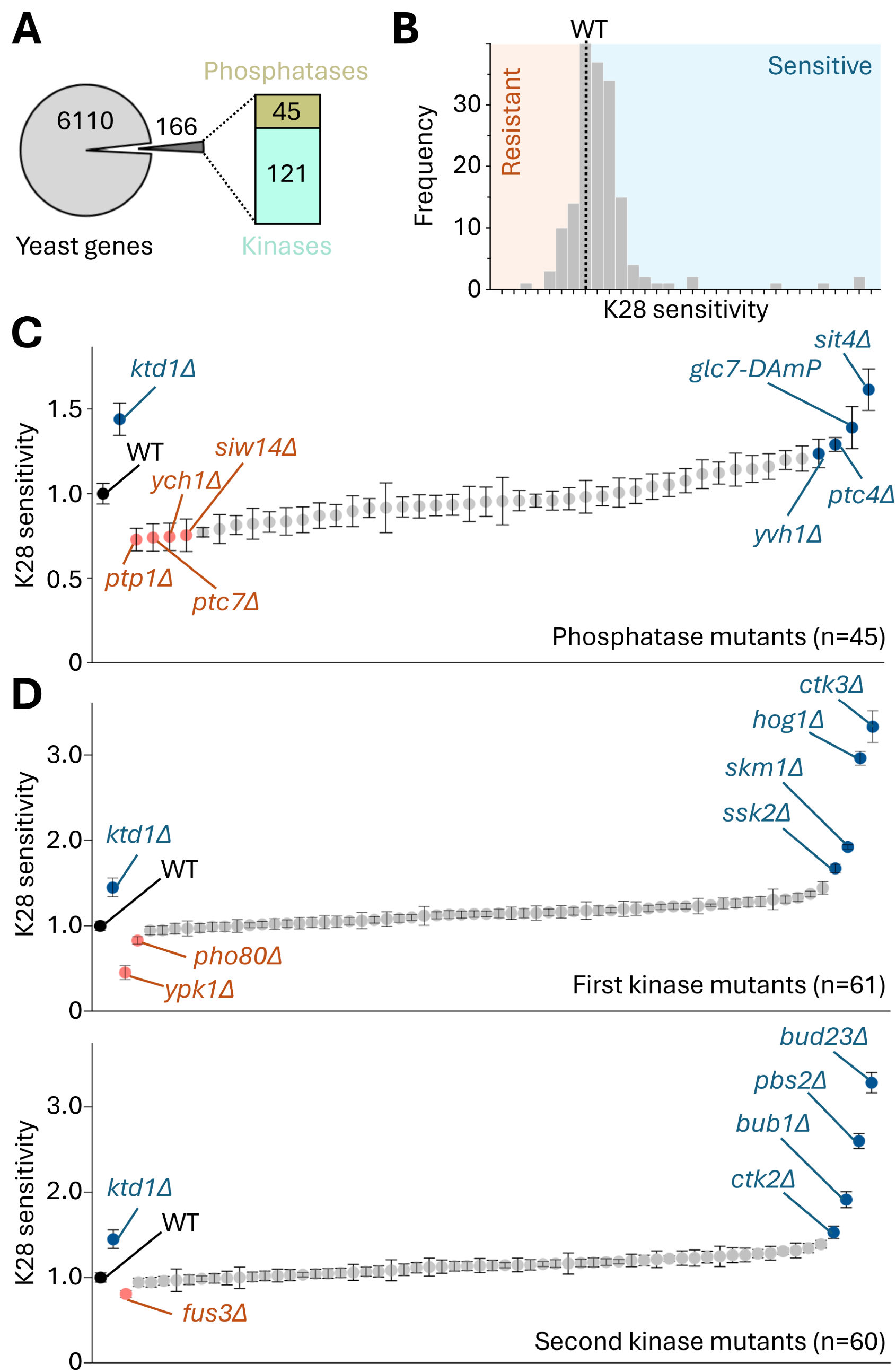
A screen for phospho-regulators of the K28 defence pathway. **A)** Mutants of 166 genes related to phosphorylation (45 phosphatase and 121 kinases), were prioritised for screening for K28 sensitivity. **B)** Each mutant was tested for K28 sensitivity (n=4) using optimised assay conditions and analysis. The distribution of phenotypes is depicted compared to wild type (WT, dotted line) and frequency of resistant (red) and sensitive phenotypes. **C - D)** K28 sensitivity of each mutant is plotted from the list of phosphatases **(C)** and kinases **(D)**, with high (blue) and low (red) scoring mutants labelled.

To validate these hits from the screen, we next performed liquid-based K28 sensitivity of the top four phosphatase and eight kinase mutants (**Figure 6A - 6C**). We note that some of these mutants have growth defects when compared to wild-type cells (**Sup. Figure S4**). However, these mutants also exhibit a hypersensitivity to K28 exposure based on liquid growth cultures compared to heat cured controls. That said, we do note that *bub1Δ* and *ptc4Δ* mutants, although hypersensitive to K28, show more subtle phenotypes than other high scoring mutants from the halo assay screen. String pathway analysis revealed several associations from the high-scoring mutants (**Figure 6D – 6E**). The most obvious being identification the Hog pathway (Saito and Posas, 2012), with the MAPKKK Ssk2, the MAPKK Pbs2, and the MAPK Hog1 itself being associated with K28 sensitivity, through a well-recognised signalling pathway. We also note whilst MAPKKKs Ste11 mutants were sensitive (1.3 ± 0.1 halo distance) the functionally redundant Ssk22 was not (1.02 ± 0.06). Similarly, *cla4Δ* mutants were hypersensitive to K28 but *ste20Δ* were not (1.0 ± 0.07). This highlights that significant pathways, but perhaps only specific arms, are require for K28 toxin defence, mimicking other stress response pathways. In combination, this provides a useful resource for all phosphorylation genes that are involved in K28 sensitivity.

**Figure 6:**
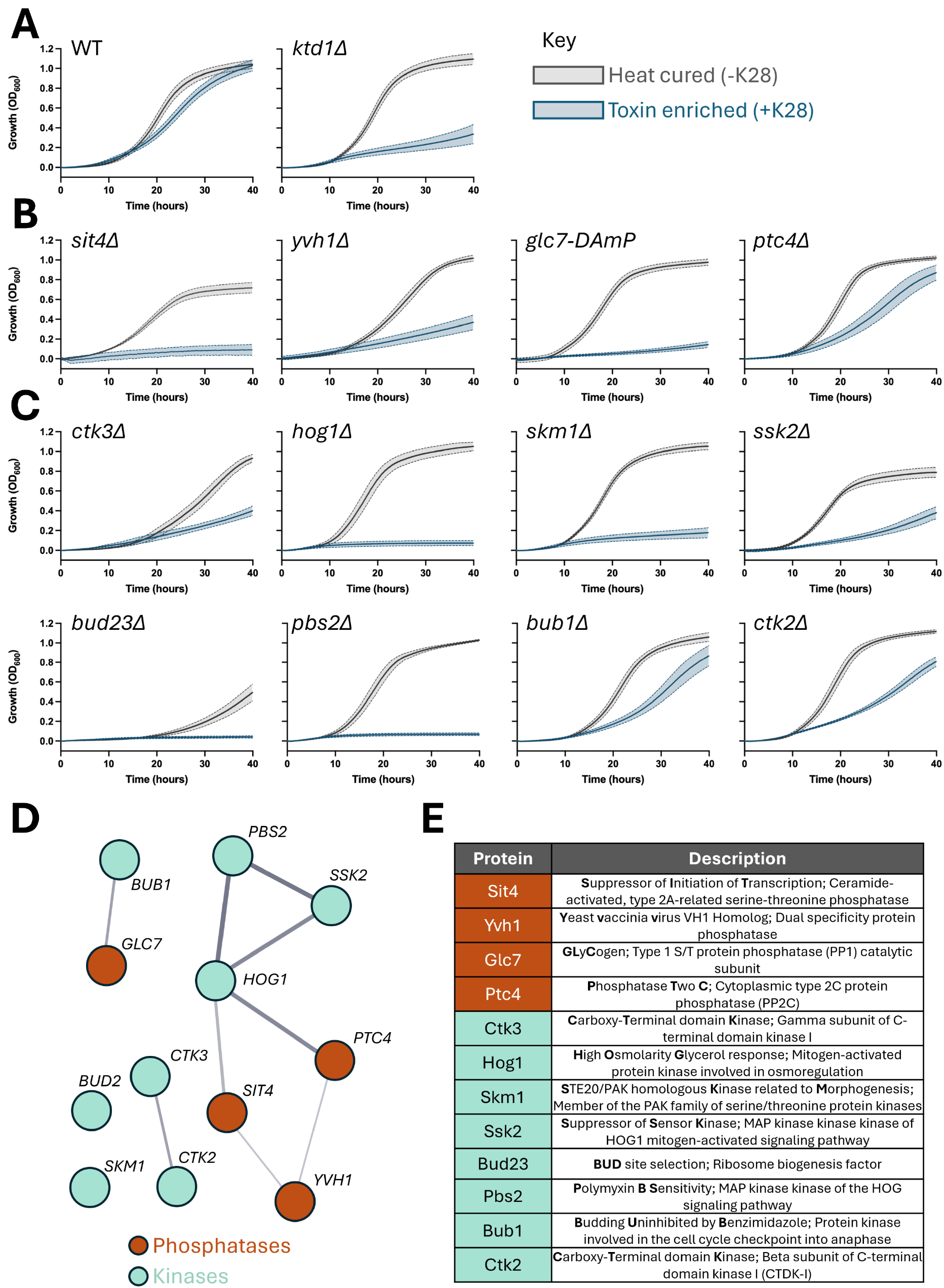
Analyses of notably K28 sensitive phospho-regulator mutants. **A - C)** Liquid based assays in the presence of K28 enriched media (blue) or a heat cured toxin control (grey) are depicted for control strains **(A)**, phosphatase mutants **(B)** and kinase mutants **(C). D- E)** Top hits from the screen (phosphatases, red; kinases, teal) are shown, with associations from STRING analysis **(D)** and molecular function **(E)** indicated.

### Hog1 and Glc7 implicated in Ktd1 mediated defence against K28

The screen tested for any phosphorylation genes involved in K28 sensitivity. We next set out to prioritise any phosphatases / kinases that might explain the phospho-regulation of Ktd1 required for K28 defence. We previously created a system for K28 sensitive assays that select for tester strains harbouring plasmids. For this, a uracil-prototrophic K28 secretor strain was engineered and used in media lacking uracil with a lawn of cells expressing GFP-tagged Ktd1 on a uracil plasmid (Laidlaw et al., 2025). To expand this toolkit, we generated a leucine-prototrophic K28 hypersecretor strain that effectively kills both wild-type and *ktd1*Δ cells (**Sup. Figure S5**), and a leucine-selectable plasmid expressing mScarlet3-tagged Ktd1 (mS3-Ktd1). Expression from the *CUP1* promoter showed mS3-Ktd1 localises correctly to the vacuolar membrane and endolysosomal compartments (**Figure 7A**). We did note that copper could be titrated to elevate mS3-Ktd1 levels, but increasing copper beyond ∼10 µM resulted in mS3-Ktd1 mislocalising to the ER, marked by Tna1-GFP. Expression of mS3-Ktd1 from a plasmid in a wild-type background with endogenous *KTD1* provides extra levels of K28 defence and functionally complements *ktd1Δ* mutants (**Figure 7B**). Comparing media supplemented with 100 µM copper to basal levels revealed no significant increase in K28 defence, suggesting that a functional Ktd1 population traffics to the endolysosomal system against the backdrop of steady-state ER mis-localisation.

**Figure 7:**
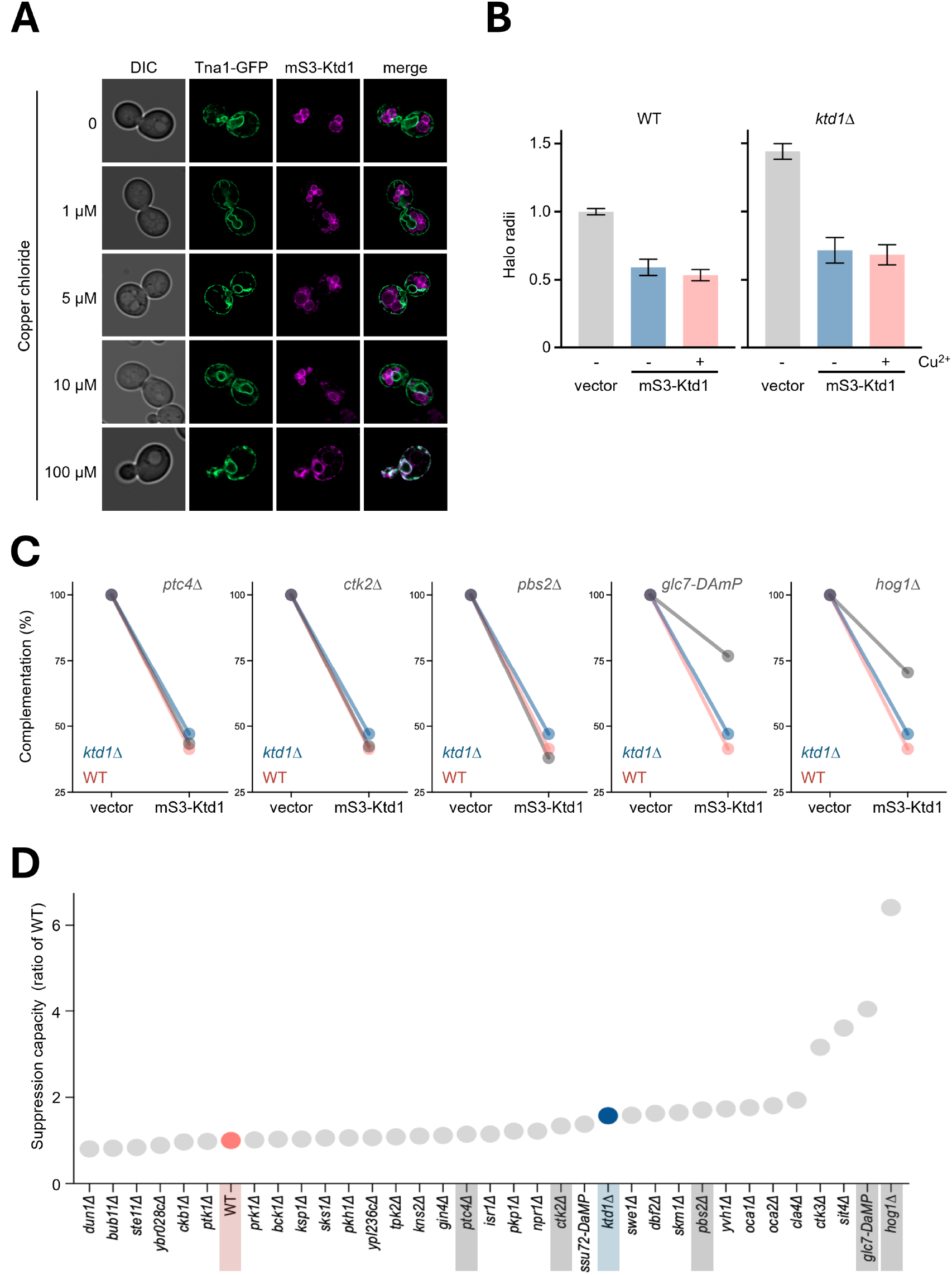
mScarlet3-Ktd1 successfully rescues sensitive phenotypes. **A)** Airyscan2 micrographs comparing mScarlet3-Ktd1 co-localisation with the ER marker Tna1-GFP upon incubation with indicated concentrations of copper chloride. **B)** Halo distances of control strains transformed with an empty vector or mScarlet3-Ktd1 (mS3-Ktd1) plasmid. Assays were performed in standard media lacking additional copper (-) or in the presence of 100µm copper chloride. **C - D)** Suppression phenotypes of indicated mutants transformed with mScarlet-Ktd1 expression plasmid, presented as a comparison with plasmid controls **(C)** or as a normalised score against wild-type rescue levels **(D)**.

As previously noted, over-expression of Ktd1 increases K28 defence in wild-type cells and rescues *ktd1Δ* mutants effectively, which was used as a visual benchmark for the rescue potential of different phosphorylation mutants (**Figure 7C**). This suppression screen was effective, with some mutants being rescued to wild-type and *ktd1Δ* levels, such as *pbs2Δ* mutants, whilst others like *glc7-DAmP* and *hog1Δ* remaining hypersensitive to toxin despite additional *KTD1* expression. To compare rescue, we normalised the suppression capacity to wild-type cells, showing *hog1Δ, glc7-DAmP, sit4Δ*, and *ctk3Δ* had most inefficient rescue by mS3-Ktd1 over-expression (**Figure 7D**). This downsteam screen indicates most hits from the K28 sensitivity screen can be rescued to near wild-type levels upon expression of mS3-Ktd1, suggesting that these enzymes act upstream of Ktd1 or via a parallel mechanism. Conversely, kinases Hog1 and Ctk3, and phosphatases Glc7 and Sit4 might be required for Ktd1 function or act downstream of Ktd1 in the K28 defence pathway.

As an initial exploration of these candidates, we assessed GFP tagged Glc7 following exposure to K28 toxin. As previously documented, GFP-Glc7 localises to multiple cellular locations depending on the cell cycle (Paine et al., 2023). We did not observe obvious differences in localisation upon exposure to K28 for 1 hour that were cell cycle specific, but we did observe a higher frequency of larger punctate GFP-Glc7 structures (**Figure 8A - 8B**). The nature of these and whether they correlate with Ktd1 activity is unknown, but there were instances when such foci were close to vacuoles labelled with FM4-64. We did not detect aggregation or other mis-localisation phenotypes of other GFP-tagged phosphorylation enzymes implicated in K28 defence, such as Ptc4, Skm1, Cla4 and Yvh1 (**Sup. Figure S6**). In response to osmotic stress, Hog1 is rapidly phosphorylated and translocated to the nucleus within 5 minutes, with dephosphorylation and return to the cytoplasmic localisation observed after 30 minutes (Bicknell et al., 2010; Reiser et al., 1999). We confirmed GFP-Hog1 behaviour by exposing cells to 0.4 M NaCl and imaging cells after 5- and 30-minute time points to show the same kinetics to stress (**Figure 8C**). Next, we exposed GFP-Hog1 cells to K28 but did not observe any obvious changes in localisation or levels (**Figure 8D**). Based on these two examples, we conclude that the effects of phosphorylation enzymes on Ktd1-mediated K28 defence are likely transient.

**Figure 8:**
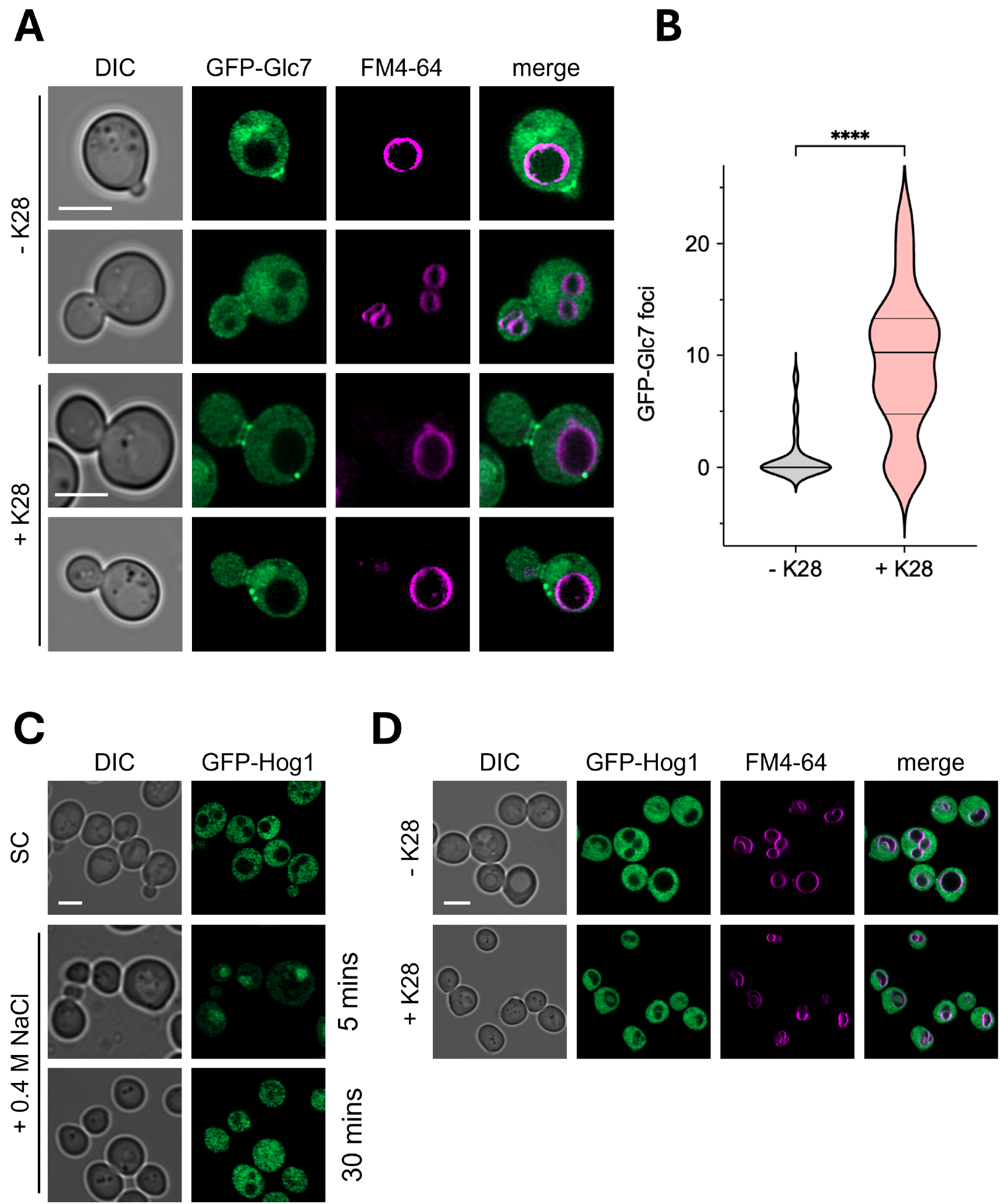
K28 exposure effects can be quantified by confocal microscopy. **A)** Airyscan2 micrographs showing the effect of adding K28 to GFP-Glc7 cells, with FM4-64 acting as a vacuolar membrane stain. **B)** Quantification of the number of foci in for GFP-Glc7 micrographs as in **(A). C)** GFP-Hog1 cells shown in synthetic complete media, compared to 0.4M NaCl solution at 5 minutes and 30-minute intervals. **D)** GFP-Hog1 cells in the absence and presence of K28 toxin, again with FM4-64. Scale bars, 5 µM.

## DISCUSSION

In this study, we performed a comprehensive functional and structural characterisation of the *S. cerevisiae* Dup240 family, establishing how sequence divergence and post-translational modifications govern their spatial distribution and role in toxin defence. Although the Dup240 family shares high sequence homology across its members, Ktd1 uniquely mediates resistance against K28 toxin. By integrating predicted structural features, sub-cellular localisation assays, and targeted phospho-mutant analysis, we demonstrate that Ktd1 function requires precise phosphorylation at conserved residues. Furthermore, our enzyme-wide array identified key upstream kinases and phosphatases, most notably Hog1 and Glc7, implicated in modulating K28 defence.

Our comparative analysis reveals that while the N-termini and initial transmembrane domains (TMDs) remain tightly conserved, key variations within the rapidly evolving luminal loop and second TMD likely dictate functional specificity across the family (Andreev et al., 2023). Ktd1 was the only family member with a significant role in K28 defence, so we sought features from the comparative analysis that might explain this specificity. Ktd1 exhibits distinct structural traits, including a predicted membrane-adjacent coiled-coil region that is absent in all other Dup240s. It is tempting to speculate that this feature might contribute to localisation of Ktd1 to the limiting membrane of the vacuole, which was not observed for any of the other five successfully localised Dup240s (**Figure 2**). In addition to a predicted structural region close to the membrane that differs from the canonical Dup240 family, Ktd1 also localised uniquely to the vacuolar membrane, whereas other Dup240 members that we could visualise were localised either to the ER membrane or the lumen of the vacuole. Ktd1 and/or the closely related Uip3 were identified in complex with the vacuolar V-ATPase (Wang et al., 2023), which may promote their secretory or post-Golgi trafficking, but additional regulation must dictate the retention of Ktd1 in endolysosomal/vacuolar membranes when Uip3 is sorted into the lumen of the vacuole. We have previously shown that the small luminal loop region is important for Ktd1 function (Andreev et al., 2023), and Ktd1 is one of several Dup240s that contain a luminal cysteine residue, which may facilitate specific protein-protein or protein-lipid interactions. Uip3, which does not contribute to K28 defence, lacks this cysteine.

This structural variation translates directly into distinct sub-cellular compartmentalisation. Unlike earlier reports suggesting plasma membrane targeting via C-terminal fluorescent tagging (Poirey et al., 2002), our functional N-terminal GFP constructs, and improved microscopy capabilities, revealed that Dup240 members partition exclusively between the endoplasmic reticulum (ER) and the endolysosomal system. While Dfp3 and Uip3 target the vacuolar lumen, Mst27, Prm8, and Prm9 localise to the ER membrane in a striking punctate pattern. This distinct microdomain organisation contrasts with uniform ER markers like translocon subunits (Prinz et al., 2000). Indeed, although we believe that cycling of Ktd1 between endolysosomal compartments is important for its function in K28 defence, we also note its localisation at the vacuolar membrane is also punctate. This could be a feature of the Dup240 family, even when localised to distinct membranes. Indeed, it may be that non-Ktd1 Dup240 proteins might regulate membrane domain architecture or lipid organisation at the ER, analogous to the role of Ktd1 within the endolysosomal network.

Another observation was that Ktd1 was predicted to be post-translationally regulated at unique sites not predicted even for closely related family members. Our bioinformatic and phospho-ablative analyses confirm that Ktd1 activity relies on two distinct serine/threonine phosphorylation clusters (Cluster 1: T3, S5, T8; Cluster 2: S170, S174, S176, S182, S185). We also note the unique predicted structural element in Ktd1 and absent from all other Dup240s (**Supp figure S1D**) is adjacent to the modifications of cluster 2. Mutating either cluster of Ktd1 to alanine completely abolished its capacity to rescue *ktd1Δ* hypersensitivity, establishing phosphorylation as an essential regulator for K28 defence (**Figure 4**). Whole-cell proteomic datasets report a higher density of modified residues on Ktd1 compared to other family members (Lanz et al., 2021; Swaney et al., 2013; Zhou et al., 2021). While this may reflect true biological differences, such as Dup240 specific regulation, we also acknowledge higher steady-state abundance, for example of Ktd1 compared to Dfp3 and Uip3 that are mainly degraded in the vacuole, might explain documented modification. In this sense, the computationally driven predictions we outline provide an extra layer of modelling to help understand differences and similarities across Dup240s. Differences in documented PTMs across Dup240s may also stem from technical detection biases during mass spectrometry profiling of specific peptides (Cunningham et al., 2012). Nevertheless, the recent discovery of a phosphorylated tyrosine (Y134) under acute stress conditions (Leutert et al., 2023) further highlights the complex, multi-layered phospho-regulation governing Ktd1. It is plausible that other Dup240 members employ analogous phosphorylation switches to respond to specific environmental stresses, operating at different cellular membranes. Beyond this, it also known that Ktd1 is ubiquitinated (Back et al., 2019; Hitchcock et al., 2003; Kolawa et al., 2013), so all Dup240 proteins might utilise a range of post-translational regulation.

Our high-throughput screen expanded the known network of phosphorylation enzymes modulating K28 sensitivity, identifying 26 novel candidates alongside several previously established enzymes (Carroll et al., 2009). The identification of the high-osmolarity glycerol (HOG) pathway, comprising Ssk2, Pbs2, and Hog1, aligns with its documented role in protecting against other killer toxins such as K1 and K2 (Gier et al., 2019; Pagé et al., 2003; Servienė et al., 2012). That said, K1 and K2 are pore forming toxins, that trigger a breakdown of the electrochemical gradient and loss of internal turgor pressure that mimics severe osmotic stress (Ahmed et al., 1999; Martinac et al., 1990; Orentaite et al., 2016; Tamás et al., 2000). Therefore, the rapid triggering of the HOG pathway might be expected. The mode of action of K28 triggering cell cycle arrest and death following toxin internalisation is thought to be different (Reiter et al., 2005; Schmitt et al., 1996), so the role of HOG might help understand specific toxin modes of cell death. However, our genetic suppression screen revealed that overexpressing *KTD1* rescued *pbs2Δ* cells but failed to restore resistance in *hog1Δ* (**Figure 7**). This indicates that Hog1 may either act downstream of Ktd1 or is directly required for its functional activation. The fact that upstream MAPKK / MAPKKK enzymes can be suppressed by over-expression of mS3-Ktd1 may be explained by small pools of Hog1 being active and sufficient in the *pbs2*Δ / *ssk2*Δ mutant background, or a more complex cross-talk mechanism with the HOG pathway. Our suppression screen also suggests Glc7 might be required for Ktd1 functional regulation. This is supported by the observation that GFP-Glc7 localises to cytoplasmic puncta following addition of K28 (**Figure 8**), but how this might correlate with Ktd1 regulation remains unclear. Live-cell imaging of GFP-Hog1 did not reveal major spatial translocations or steady-state alterations following addition of K28, suggesting its role may be transient amidst the large background of additional functions (Hohmann, 2002; Nadal and Posas, 2022).

The other screen hits will also serve as useful models for understanding potential modes of K28 defence. For example, Glc7 and Sit4 are both known to have roles in the endolysosomal system that might integrate with Ktd1-mediated defence (Bowman et al., 2022; Como and Arndt, 1996; Martins et al., 2024; Merhi and André, 2012). For example, *tda1Δ* mutants are hypersensitive (halo distance 1.28 ± 0.05); Tda1 is a kinase that regulates glucose signalling (Kaps et al., 2015; Nonaka et al., 2025) and is required for K28 defence, which might integrate with glucose-mediated control of endocytic internalisation or recycling (Laidlaw et al., 2021; Laidlaw et al., 2022) necessary for K28 entry or correct Ktd1 localisation. Other examples of phosphorylation enzymes worthy of mechanistic dissection in future studies are Env7, a kinase that localises to the vacuole and Ybr028c, a putative kinase required for efficient defence (Manandhar et al., 2013; Ricarte et al., 2011). These datasets and resources will aid future hypothesis generation and a mechanistic understanding of how eukaryotic cells tolerate the internalisation and killing action of AB toxins.

## METHODS

### Reagents

Plasmids and yeast strains used in this study are detailed in Supplemental Table T2.

### Cell culture

A combination of rich and minimal media was used for yeast cultures depending on requirements. Rich media (YPD CCM1010, Formedium]) contains yeast extract (1%); peptone (2%); adenine sulfate (0.004%); dextrose/glucose (2%). Minimal synthetic complete (SC) media contains glucose (2%), yeast nitrogen base without amino acids (CYN0410, Formedium), and a cocktail of appropriate amino acids and bases (i.e. DSCK1000, Formedium). 2% agar was added prior to autoclaving when making plates. Early-mid log phase cultures were grown prior to experiments following overnight incubation at 30°C with shaking.

### Sequence alignments

The ‘MUSCLE Default’ substitution matrix was used for sequential alignments of Dup240 proteins, as per the default set-up using the MegAlign Pro software (Lasergene). Phylogenetic analysis and tree creation was created through Maximal Likelihood: RAxML, with 100 iterations of bootstrap analysis. Sequence identity percentages were created using an uncorrected pairwise distance metric with pairwise gap removal.

### Precited structural visualisations

AlphaFold3 (Abramson et al., 2024) was used to predict the structures of the Dup240 proteins and create a Dup240 consensus model. The structure with the highest-ranking score (out of the five generated) was then visualised in PyMOL Molecular Graphics System (Version 3.0, Schrödinger). Rainbow spectra were applied based on the B-factor values found in the .cif file. Any alignments were performed relative to Ktd1, either the entire structure or its individual components.

### K28 halo assays

OD_600nm_ measurements were used to normalise lawn densities for assessment strains cultured to saturation through overnight incubation. Plates used for halo assays were synthetic complete containing 11.2% (v/v) phosphate citrate buffer pH 4.7. Normalised lawns were formed through a 3-minute exposure to 1500μL of OD_600nm_ = 0.025 of each assessment strain, before excess was poured off. Plates were left in the laminar flow hood for approximately 40 minutes to dry. Upon the dried lawn, between four and seven 7.5μL spots of the secretor cell culture (a *ski2Δ/ski2-2* diploid strain infected with the M28 virus grown to OD_600nm_ ≈ 1 concentrated to OD_600nm_ ≈ 10) were added. Measurements were taken after 48 hours of growth at room temperature, either by using ImageJ or an automated Python pipeline (Pembery et al., 2026).

### Purification of K28-enriched media

K28 secretor cells (*ski2Δ/ski2-2* diploid strain infected with the M28 virus) and a heat-cured negative control strain were grown in SC media pH = 4.7 for 40 - 45 hours at 23°C and 70 RPM shaking. The media enriched with toxin was then filtered with 0.22µm filter into a clean and sterile glass conical flask. 12mL of the toxin enriched media is then supplemented with 3ml of 5x concentrated SC media nutrient solution. Following filtration, the enriched media was used, either for liquid-based growth assays or for incubation with cells prior to microscopy.

### Liquid K28 sensitivity growth assay

A 96 well microtiter plate was prepared with wells containing 200µl of either K28 toxin containing media or heat cured control cultures. Strains for assessment were cultured to saturation overnight, then 1.5µl added to a minimum of 6 wells per experiment for both conditions. Certain wells containing both K28 and control media but lacking yeast inoculation were routinely prepared to control for any contamination growth. Plates were then incubated in an Alto Plate Reader (Cerillo) at 25°C, and OD_600nm_ absorbance measurements were taken every 3 minutes for approximately 27 - 40 hours I n. Raw data were plotted as line graphs using GraphPad Prism v10., with shaded regions showing standard deviation across experiments. Representative plots (from at least n = 3 biological replicates) are shown with on-the-day controls plotted for comparisons.

### Confocal microscopy

For confocal microscopy, yeast cells were first cultured to mid-log phase prior to harvesting and preparation for imaging. Microscopy was performed on a Zeiss 980 laser-scanning confocal microscope equipped with an Airyscan2 detector and a 63x Plan-Apochromat objective lens (1.4 Numerical Aperture, Carl Zeiss AG, Oberkochen, Germany) imaged using immersion oil. GFP-tagged proteins were excited with 488nm wavelength excitation, (500 - 550 nm wavelength emission). FM4-64 (*N*-(3-Triethylammoniumpropyl)-4-(6-(4-(Diethylamino) Phenyl) Hexatrienyl) Pyridinium Dibromide) dye (Thermo Fisher Scientific) was used to label vacuolar membranes by incubating mid-log phase cultures with the dye in YPD at 30°C for 30 minutes, followed by washing and resuspension in SC medium for 1 hour. FM4-64 and the red monomeric fluorescent protein mScarlet3 used to visualize Ktd1 using 561nm wavelength laser excitation (580 - 620nm wavelength emission). SC media containing 50 µM CMAC (7-amino-4-chloromethylcoumarin) dye (Thermo Fisher Scientific) was used to label the vacuolar lumen, using 405 nm wavelength laser excitation (422-497 nm emission). Images were acquired with optimized settings, followed by visualization and analysis using Zen blue and Fiji/ImageJ software.

## Supporting information

Raw data for K28 genetic screen

Reagent table

## ACKNOWLEDGMENTS

We would like to thank staff at the York Bioscience Technology Facility for technical assistance. This research was supported by a Sir Henry Dale Research Fellowship from the Wellcome Trust and the Royal Society 204636/Z/16/Z (CM) and the Engineering and Physical Science Research Council EP/Y000501/1 (MCL). We also thank Meru Sadhu, Padraic Heneghan, Dani Ungar, Gareth Evans for useful conversations related to the project.

