## Supplemental figures for "Ktd1 is a phospho-regulated member of the Dup240 family that mediates defence against killer toxin K28"

### SUPPLEMENTAL INFORMTAION

*List of supplemental figures and legends.*

***Supplemental Table 1: Halo assay screen results***
*This includes all halo distances averages, standard deviations and is coloured coded to match Supplemental Figure S3A*

***Supplemental Table 2: Reagents***
*List of plasmids and yeast strains used in this study*

***Supplemental Figure S1***
*Computational analysis of Dup240 family*

***Supplemental Figure S2***
*Predicted orientation of Ktd1 within the vacuolar ATPase complex*

***Supplemental Figure S3***
*Fluorescent markers of the endoplasmic reticulum*

***Supplemental Figure S4***
*K28 sensitivity and control growth comparisons*

***Supplemental Figure S5***
*Creation of a Leu$^+$ prototroph K28 secretor strain*

***Supplemental Figure S6***
*Localisation of phosphorylation enzymes following K28 treatment*

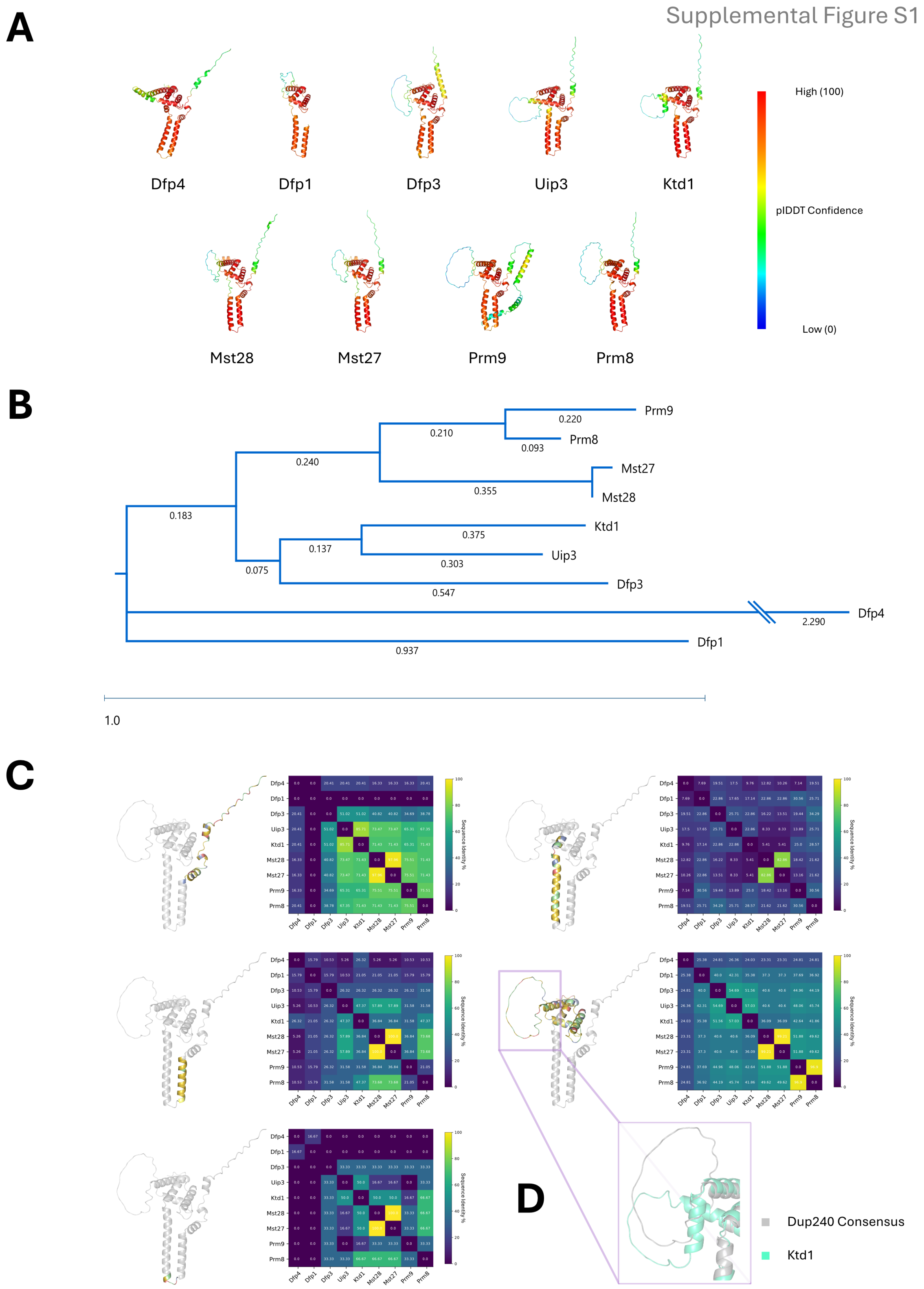


**Supplemental Figure S1: Computational analysis of Dup240 family**
**A)** AlphaFold3 predicted structures for nine of the ten Dup240s found in *S. cerevisiae* with prediction confidence shown as heat map, high (red) and low (blue). **B)** Phylogenetic tree generated from proteins in **(A)**, calculated by RAxML bootstrap analysis. Scale bar represents the evolutionary distance. **C)** Consensus structures shown in **(1B - 1F)** accompanied by heatmaps showing the sequence identity matrix between each Dup240 member. **D)** Enhanced zoom of large loop in cytoplasmic domain comparing the Dup240 consensus (grey) and the unique Ktd1 feature (blue).

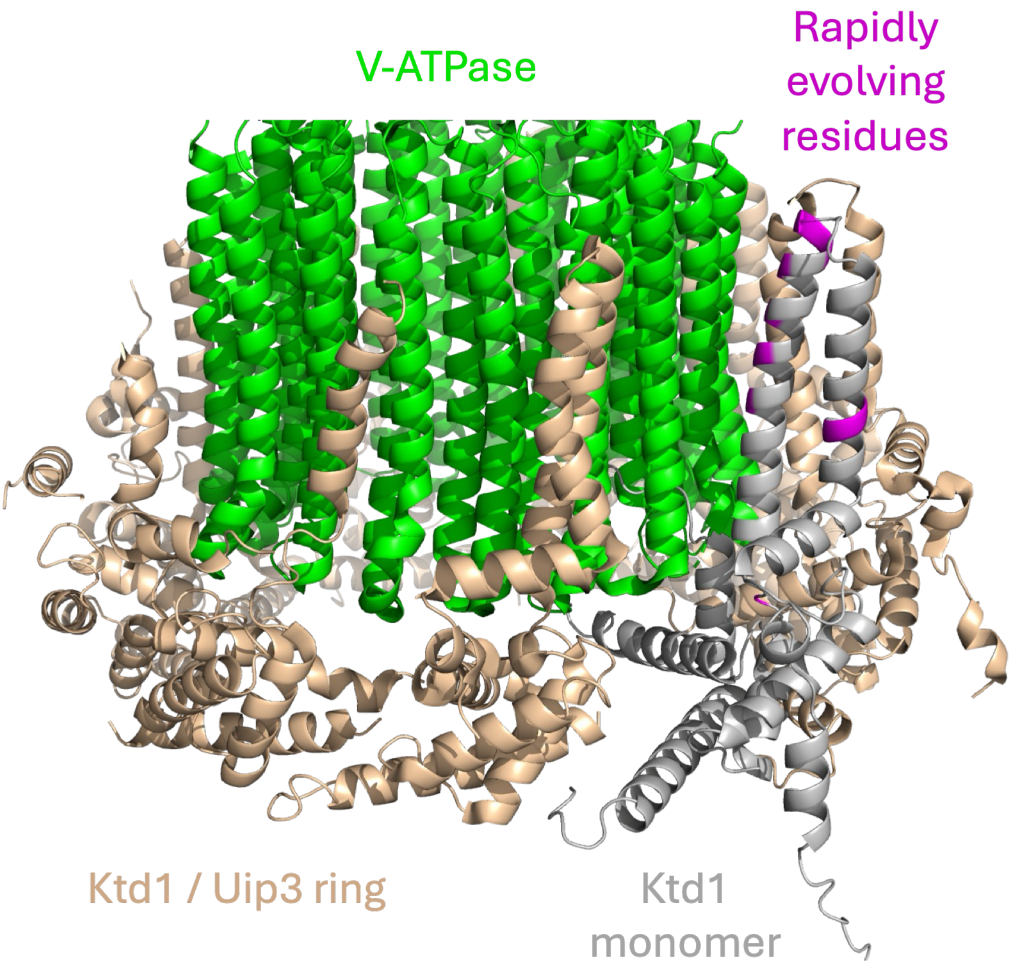


**Supplemental Figure S2: Predicted orientation of Ktd1 within the vacuolar ATPase complex.** Structure of Uip3 and Ktd1 (encoded by *YAR027W* and *YAR028W*, respectively) forming an oligomeric ring (wheat colour) in complex with the V0 c subunits (green) of the yeast V-ATPase (PDB: 8EAV; 5.7 Å resolution by single-particle cryo-EM; Wang *et al.,* PNAS 2023). A single monomer of Ktd1 is highlighted (grey), with rapid evolving residues indicated (magenta).

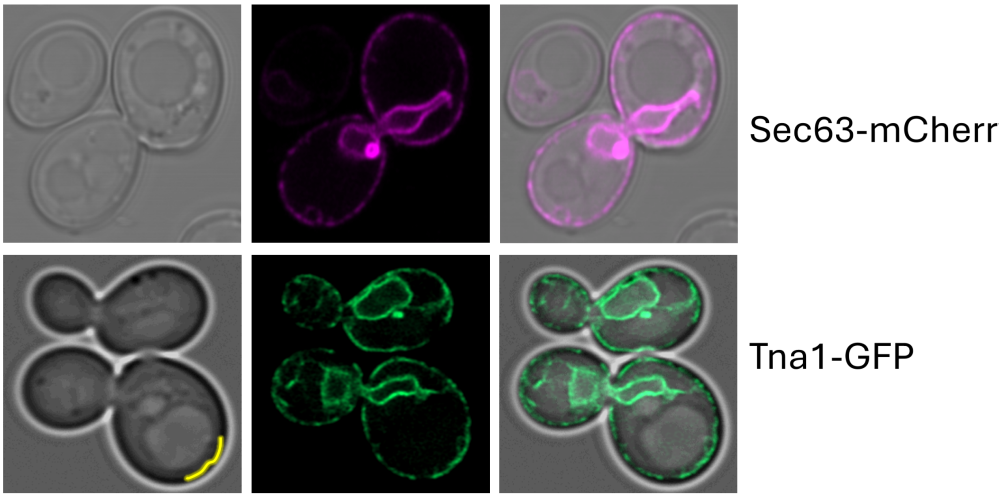


**Supplemental Figure S3: Fluorescent markers of the endoplasmic reticulum**
Airyscan2 micrographs showing Sec63-mCherry (upper) and Tna1-GP (lower). The yellow line shown in the brightfield Tna1-GFP micrograph (left) was used for the line analyses in Figure 2C.

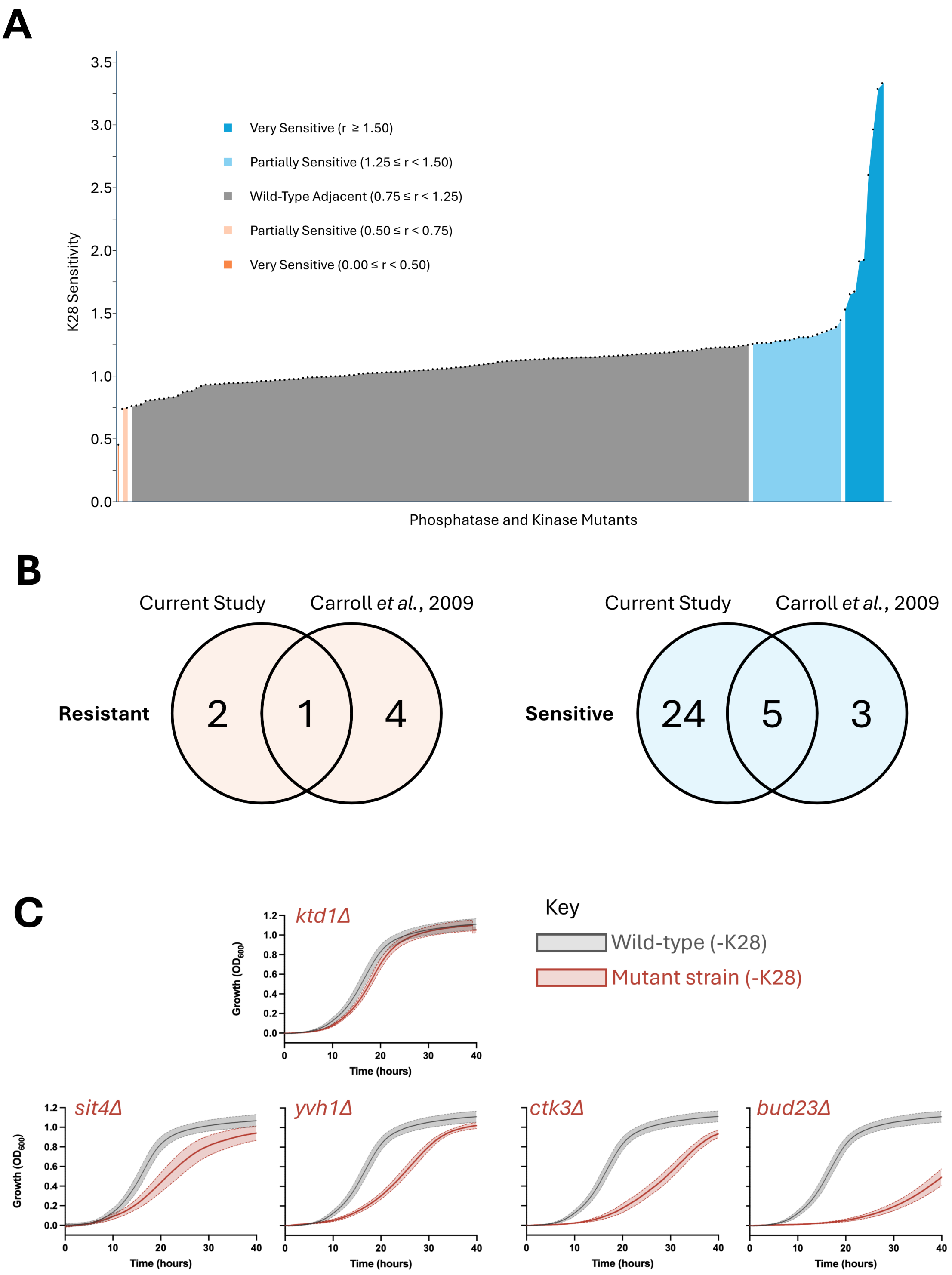


**Supplemental Figure S4: K28 sensitivity and control growth comparisons**
**A)** Colour segmented plot for all 166 genes related to phosphorylation that were screened for K28 sensitivity. Thresholding for colours was adapted from Carroll *et al.* 2009 to denote the various phenotypes upon exposure to K28 in halo assays outlined in Figures 5A - 5D. **B)** Venn diagram comparison between our current study and Carroll *et al.* 2009 genome wide K28 sensitivity screen. The intersection represents the number of mutants that both studies demonstrated have identical sensitivity phenotypes. Resistant (red) and sensitive (blue) encompasses both “very” and “partially” denoted phenotypes. **C)** Liquid growth assays of indicated mutants (red) compared to wild type controls (grey) grown under same conditions.

**A**

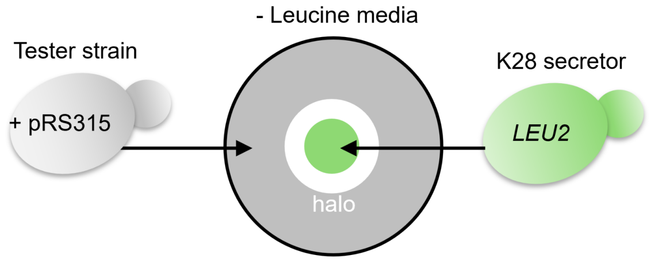


**B**

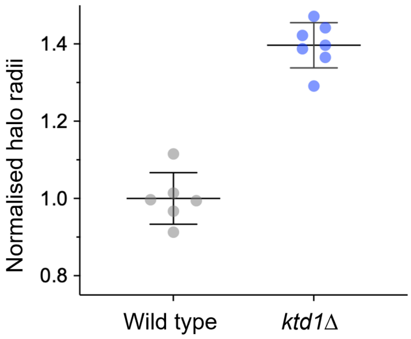


**Supplemental Figure S5: Creation of a Leu⁺ prototroph K28 secretor strain**

**A)** Schematic for a halo assay that can be performed whilst selecting for Leu$^+$ transformants based on the pRS315 backbone. To ensure efficient K28 secretion, the *LEU2* gene in M28 infected cells was repaired (see supplemental regents table for genotype information). **B)** Halo distances were measured for wild type (grey) and *ktd1Δ* (blue) cells from halo assays performed in SC media using the Leu$^+$ prototrophic secretor strain.

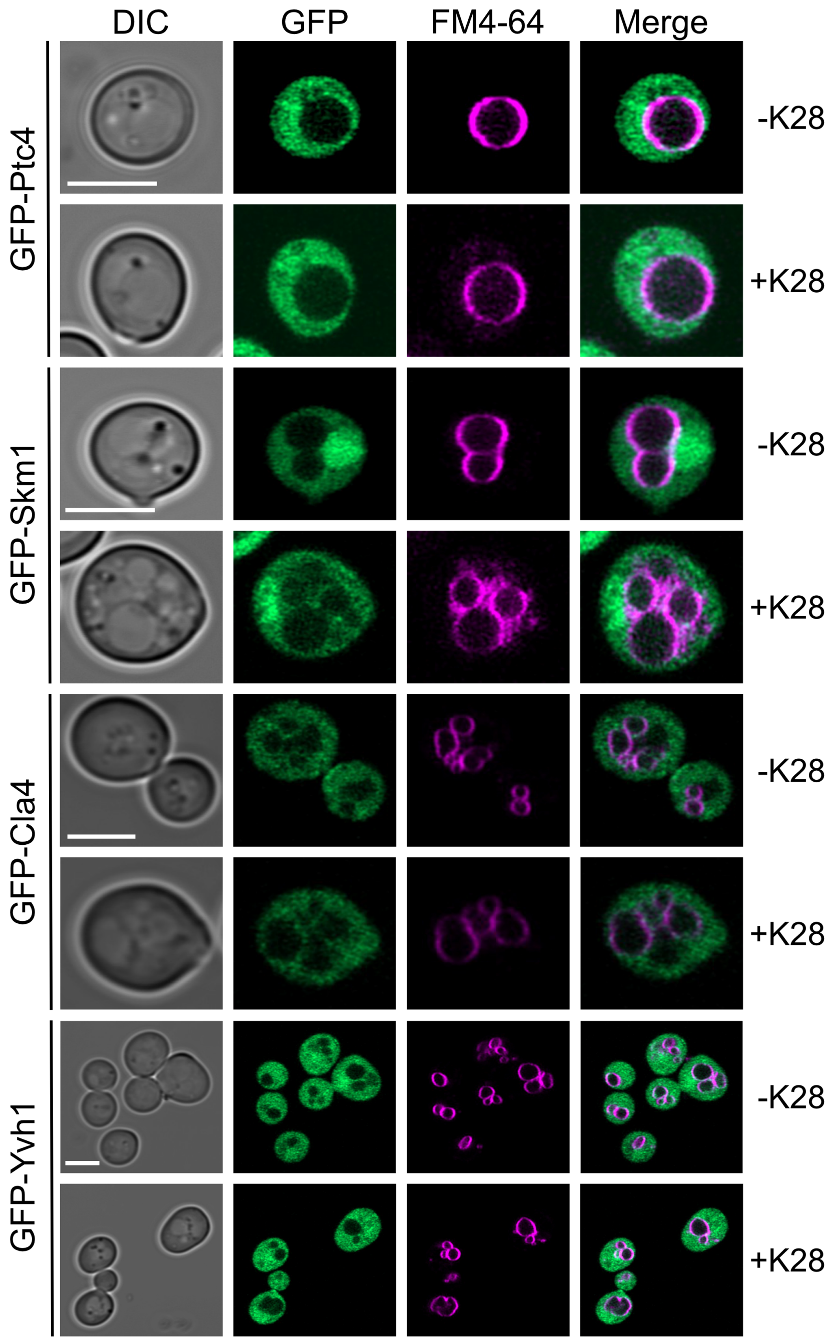


**Supplemental Figure S6: Localisation of phosphorylation enzymes following K28 treatment**
GFP strains in the absence and presence of K28, using FM4-64 as a vacuolar membrane marker. Micrographs taken using an Airyscan2 confocal microscope. Scale bar, 5 µm.
